# Visual Attention Dynamics Entrain to an Auditory Beat: The Palimpsest Paradigm

**DOI:** 10.1101/2025.08.26.672382

**Authors:** Michelle J. Spierings, W. Tecumseh Fitch

## Abstract

We investigated how rhythmic auditory cues influence visual perception of dynamically presented, partially masked words. Participants were presented with an ever-changing array of multiple overlapping, semi-transparent five-letter words, in a stimulus we dubbed a “palimpsest”. These letter arrays mostly consisted of nonsense strings, but every two seconds, two actual words briefly appeared consecutively (and typically subliminally), superimposed upon the nonsense letter array. Participants were asked to name all words that they recognized. One group of participants observed only visual stimuli, while a second group additionally received an acoustic stimulus: a drum beat that was timed to co-occur with some, but not all, of the words in the array. Participants provided with this additional synchronized auditory information were able to detect more words from the array, and the words that they recognized were typically those that coincided with the dominant acoustic beat. This shows that when visually presented words are difficult to detect, acoustic cues can improve word recognition by dynamically focusing participants’ attention upon specific moments in time. We also found that the auditory rhythm continued to affect visual perception after the sound fade fades away. A third group heard the drum beat only for the first four seconds, after which it faded away for the remaining 76 seconds of a trial. Like the continuous beat in condition two, the transient acoustic rhythm continued to influence participants’ perception of the written words, even after its cessation. These results are consistent with the hypothesis that dynamic, temporal aspects of visual perception can be strongly influenced by periodic auditory input, and that this results from the entrainment of an endogenous multi-modal attentive process. The implications for neural models of dynamic attention are discussed.

## 1. Introduction

A persistent theme in experimental psychology is the notion that the mind packages incoming stimuli into a series of “psychological moments,” at a rate of roughly ten moments per second (White, 1963). Early workers proposed two models for this phenomenon, one in which all information within a subjective moment would be summed, thus blurring or “overprinting” successive rapid events, and an alternative model in which perception is itself phasic, with peaks and troughs of attention within a single perceptual moment. Research involving rapid serial presentation of word lists argued strongly for the latter model: when multiple words were presented in rapid succession, some were perceived sharply and clearly and others were not perceived at all (Lawrence, 1971). Whether words were simply counted, or the specific words batch-reported at the end of a list presentation, made no difference, suggesting that the maximum rate of word perception of about 6 words per second represents a true perceptual “speed limit” rather than a motor limitation on counting rate. This early research suggested that visual attention is deployed dynamically and characterized by a rate of succession of moments (periodicity) and a rise and fall of attention within a moment (phase).

In a parallel research track in the musical domain, Mari Riess Jones (1976) further argued that both the period and phase of dynamic auditory attention can be entrained by periodic stimuli, such as a sequence of musical notes. Many subsequent models following from Jones’s “dynamic attending theory” share the notion of an endogenous attentional oscillator which exhibits stable, periodic persistence even after exogenous rhythmic cueing has ceased (Jones 2089; Haegens & Zion Golumbic, 2018). If so, once dynamic attention has been entrained to a given frequency and phase, it will continue to preferentially expect events in line with that phase. In the musical context, this means that once a sequence of notes establishes a clear rhythm and meter in a listener, this inferred rhythmic structure will persist in the face of (mild) counter-evidence (e.g. syncopations: events which misalign with the inferred rhythm). When counter-evidence becomes too strong (highly syncopated patterns) the endogenous oscillator should reset its phase, yielding a reinterpretation of the pattern. This prediction is upheld in experiments involving a range of mildly- to highly-syncopated rhythms (Fitch and Rosenfeld, 2007).

Despite their evident conceptual similarities, these two literatures (visual word list presentation and musical rhythm perception) have developed essentially independently. But more recent work examining cross-modal integration of auditory and visual information suggests a promising way to potentially unify the insights from these two separate sub-fields and sensory domains (Sekuler *et al*., 1997). Recent research that combines rapidly flashed images with auditory tones suggests that auditory stimulation can strongly influence visual attention: participants simultaneously exposed to a single visual stimulus and two auditory stimuli (beeps) reliably report “seeing” two stimuli (Shams *et al*., 2000). The one-way directionality of this influence (auditory-to-visual) is presumably related to the superior temporal resolution of the auditory system (Recanzone, 2003; Watkins *et al*., 2006). This and other research on cross-modal integration suggests that it should be possible to “drive” the period and phase of *visual* attention by using appropriately timed *auditory* stimulation (for reviews see Koelewijn *et al*., 2010; Spence, 2011; van Atteveldt *et al*., 2014).

In the research presented here we attempt to synthesize the insights and methodologies of these different literatures by developing a cross-modal paradigm for exploring dynamic attention and its entrainment by dynamic stimuli. We investigate auditory influences on phasic visual attention using a rapidly changing visual overlay of random semi-transparent alphabetic strings – a ‘palimpsest’- in which words occasionally appeared. When detected, words were to be spoken aloud by participants. In a baseline condition, palimpsest videos were presented silently; in an auditory beat condition they were accompanied by a periodic rhythm track in which an emphasized beat coincided with certain partially masked words, but was asynchronous with other words. We hypothesized that synchronous visual words would be detected at significantly higher rates than asynchronous words.

Further, to test the hypothesis of dynamic attending models that an endogenous dynamic attentional process can be entrained by a rhythmic stimulus (Jones, 1976; Jones and Boltz, 1989), a third condition began with a few seconds of a synchronized rhythmic beat which then faded to silence. Some hidden words remained synchronized with the previously cued temporal framework and others remained asynchronous. Dynamic attending theory would predict that after acoustic establishment of an entrained attentional rhythm, the synched words should remain more salient, even in the absence of any immediate auditory cues.

## 2. Methods

### 2.1 Subjects

A total of 36 participants took part in this study (18 males, 18 females, age 21.3 +/- 5.4). All participants were Austrians (native German speakers) and all reported that they lacked any visual or auditory impairments, or reading difficulties. All procedures were approved by the ethical committee of the University of Vienna (approval #00259), and all participants signed an informed consent prior to participation and received a five euro payment for participation.

### 2.2 Stimuli and apparatus

The stimuli were presented by a custom Python program that ran on an Apple Mac mini computer. Participants were seated 50 cm from the screen, facing it directly (display monitor frame rate: 60 Hz, resolution: 1280 × 720 pixels). The mean luminance per greyscale level of the stimuli was measured and calibrated with a Konica Minolta LS-150 Spot Luminance photometer. Gamma-correction was verified on a regular basis to ensure precise calibration throughout the experiment.

The stimulus consisted of an array of five rectangles (5 × 5 cm) presented in the middle of the display. In each rectangle, 10 semi-transparent letters overlapped each other at each given frame. The stimuli were presented at a rate of 20 frames per second (50 ms duration). Twice every 40 frames, some of the letters formed a common German word. Stimuli were constructed such that two successive words appeared every 2 seconds, with random letter assemblages intervening. These words appeared first in the lightest greyscale and increased in darkness over five frames (250 ms). The two words were presented with no overlap, but one appeared immediately after the other. For example, if the first word appeared at frame 5 and increased in darkness until frame 9, the second word would appear in frame 10 and darken until frame 14. Other letters appeared in different randomly chosen greyscales to partially mask the words and make them difficult to read (stimuli examples in •).

For one group of 12 participants, the visual stimuli were accompanied by auditory stimuli, consisting of a string of drum beats (500 ms ISI, so 2 Hz or 120 beats per minute) with a stronger beat co-occurring with one of the two words (every 2000 ms). There were three off-beats between each main beat, the total of the four beats being exactly the time between two sets of words (figure 2). By slightly shifting the start time of the beat sequence, the main beat co-occurred with the darkest frame of either the first or the second word of the visual stimuli. To provide an impression of simultaneity, the onset of the co-occurring beat was always 150 ms earlier then the appearance of the word-frame (empirically decided during pilot research). The auditory stimuli were played through headphones (Sennheiser PXC 480) at a loudness of 70 dB. The group of participants that did not receive the auditory stimuli were also requested to wear the headphones, although no sound was played.

The words used in this experiment were common German/Austrian five-letter words (mean log word frequency >1.5). They did not contain any umlauts or ß, nor where they names of people or places. Two words that occurred sequentially were matched for their mean log word frequency and the number of lexical neighbours.

### 2.3 Experimental design

At the start of the experiment, each participant was trained with five short training videos. Each video contained three sets of two words and lasted a total of six seconds. Participants were asked to call out the word that they saw appearing on the screen and their responses were recorded. After each training video, they received feedback from the experimenter on whether the words they called out actually appeared on the screen.

Participants were then informed that they would be seeing longer videos, still had to call out the words they recognized but would not receive any feedback. They were exposed to two sequential test sessions of 80 seconds each. In one test session, the beat co-occurred with the first of two words, while in the other session the beat co-occurred with the second word. This order was counter-balanced over participants.

The vocal responses of the participants were recorded during the entire experiment (ZOOM H4next microphone) and were later transcribed.

### 2.4 Analysis

The participant responses were calculated as fraction of On-Beat and Off-Beat words that were correctly recognized, as well as the proportion of words that were called incorrectly. One-way ANOVAs were used to test for significant differences between the fractions of On-Beat and Off-beat recognized words. Post-hoc Tukey tests were used to determine differences between conditions. All statistics were done in R (version 1.0.136).

Because pilot participants found the task very challenging and initially reported perceiving no words, the training session was arranged with graded difficulty – the first four training sessions involved frequent words and the fifth and final session less frequent words. This allowed participants to build confidence before beginning the test session

## 3. Results

### 3.1 Training

The five training sessions that participants were exposed to all consisted of 3 word-pairs, creating a total of 6 words per training session. Participants in the Beat condition correctly identified more words in the 3rd training session, compared to the other two conditions (mean B=2.6, mean NB=1.2, mean FB=1.2, *p*=0.03) All participants managed to correctly identify 2 to 3 words in the 4^th^ training session regardless of the sound condition they were in (figure 1). In the 5^th^ training session, in general less words were identified than in the 4^th^ session and participants from the No Beat condition performed worse than the Beat or Fading Beat conditions (mean B=2, mean NB=0.4, mean FB=1.4, *p*=0.01). This may be explained by the fact that the words in this session 5 were less frequently occurring words.

**Fig. 1.**
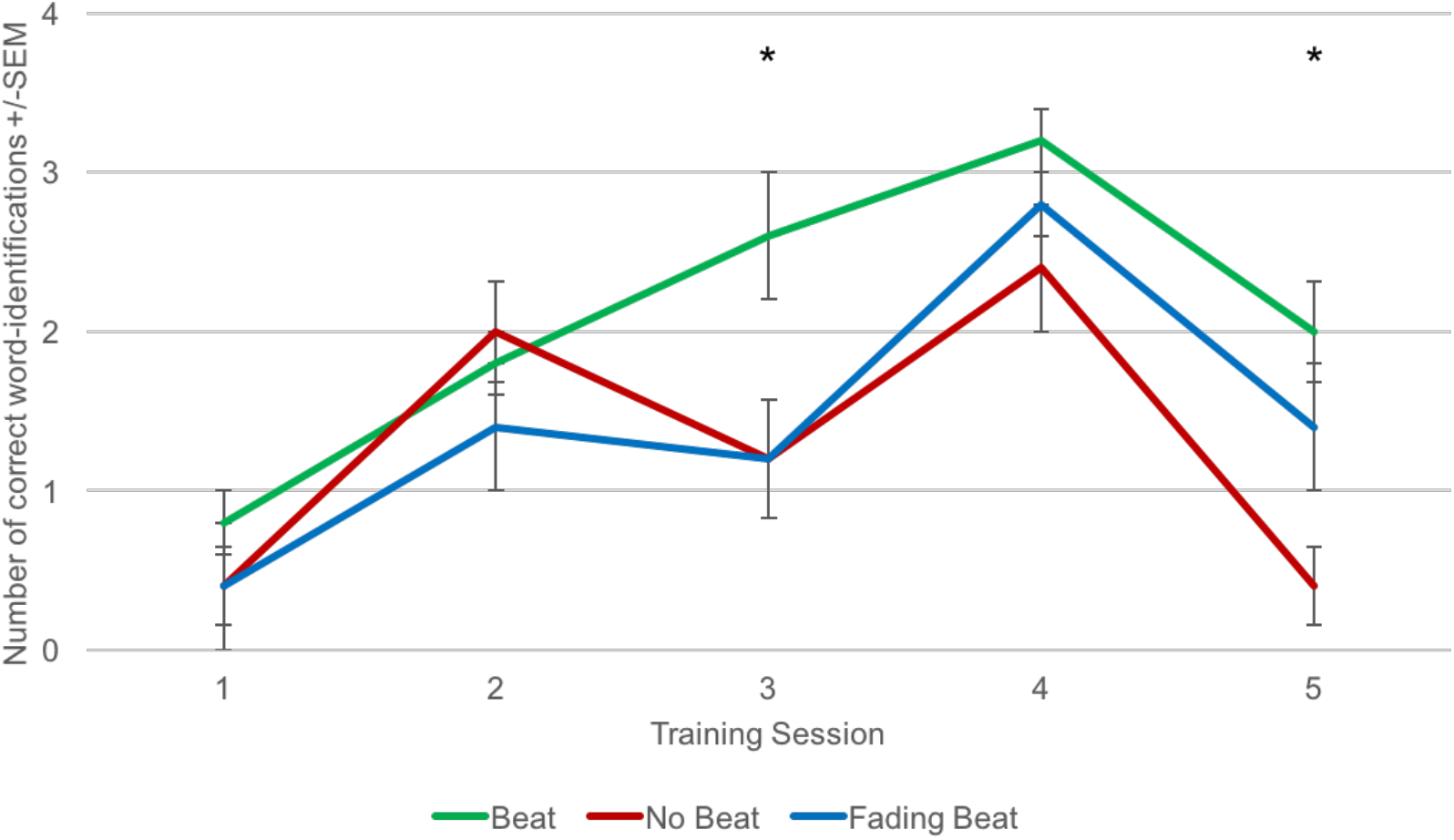
The average number of words that were identified correctly during the training sessions (1-5). In all three sound-conditions, participants recognized around 3 out of 6 words in the fourth training session. The words used in training session 5 were less frequent and therefore more difficult to identify.

### 3.2 Test

Overall, participants identified twice as many words that co-occurred with the beat than words that were either before or after the beat (*M* on-beat: 0.41, *M* off-beat: 0.21, *p*<.001, figure 2). The fraction of correctly identified words was similar for off-beat words as it was for all words in the No Beat condition (*M* off-beat: 0.21, *M* no-beat: 0.34, *p*=.24). Only in the Fading Beat condition, participants identified less words when the beat was on the second word instead of the first (*M* first-beat: 0.23, *M* second-beat: 0.15, *p*=.004). In both the Beat and the No Beat conditions, there was no effect of whether the beat was on the first or the second word (Beat, *p*=.69, no-beat, *p*=0.85). The fraction of wrongly identified words was similar amongst all conditions (mean B = 0.12 ± 0.09; mean NB = 0.08 ± 0.1; mean FB = 0.12 ± 0.08). There was no effect of sex or age of the participants on their performance in this task (*p*=.92, *p*=.86).

**Fig. 2.**
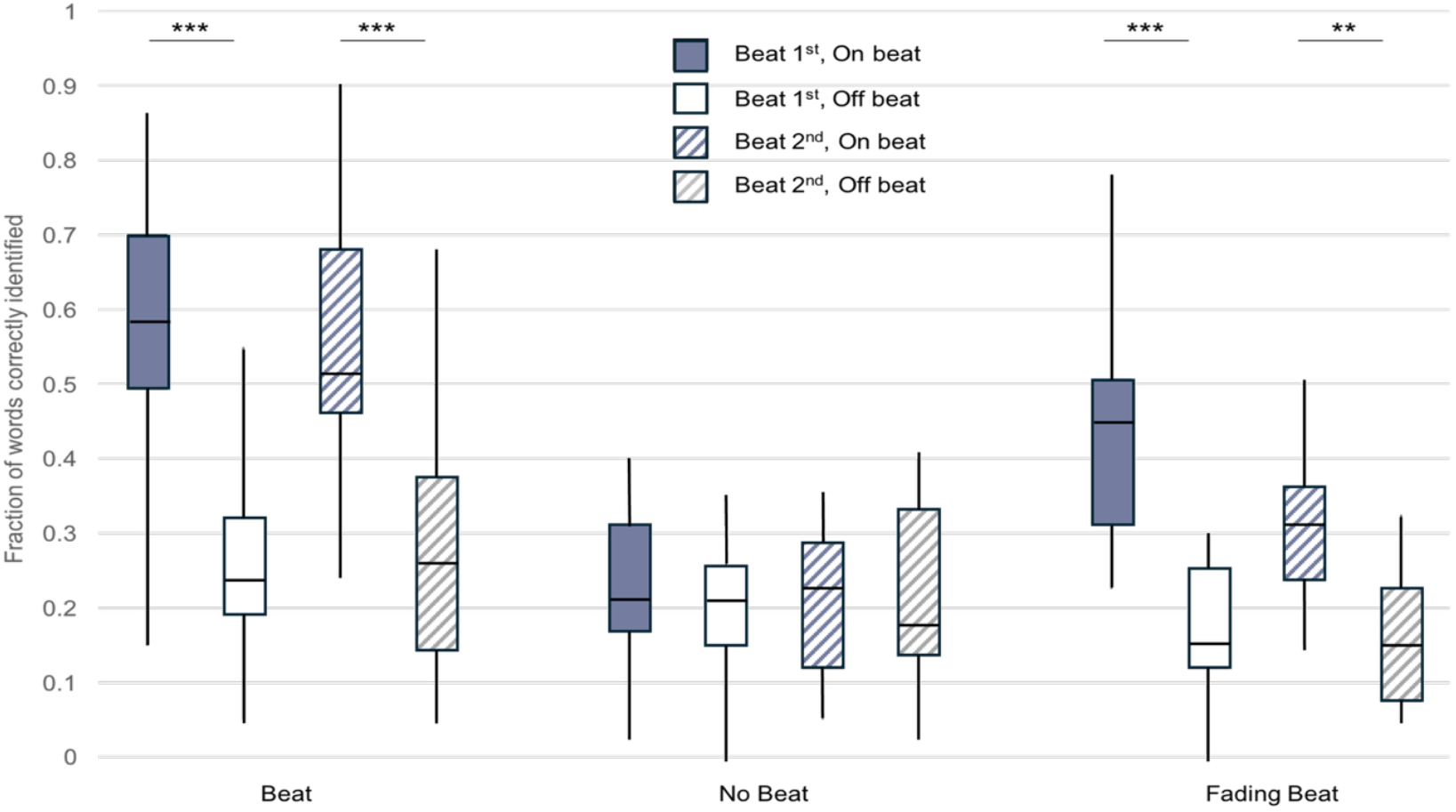
Fractions of words that were correctly identified by the participants in the three sound conditions, Beat, No Beat and Fading Beat. The grey bars show their identification words that were on the beat, the white bars of words that were not on the beat. Bars with a solid fill show results when the beat was on the first word, the striped bars show the results when the beat was on the 2^nd^ word.

Participants in the fading beat group performed best for the first 20 words that occurred directly after the beat disappeared (*M* on-beat:, *M* off-beat:) as compared to the other 60 words of the test. Each individual had a spike in recognition of the words at some later point in the test, which varied per individual. This spike naturally occurred when two words in a row were recognized correctly. The individuals seemed to pick up the beat again and recognized between 5 and 8 words sequentially. No pattern was detected onto when this spike occurred.

## 4. Discussion

This study demonstrates that visual recognition of words can be modulated by a concurrent auditory rhythm. Words that co-occur with the beat were detected more reliably than words that do not co-occur, indicating that the temporal structure provided by sound can guide visual attention. Importantly, this effect persisted to some extent even after the rhythmic cue stopped, suggesting that an isochronous rhythm, once entrained, continues to influence perception in the absence of ongoing stimulation.

These results extend prior work on multimodal integration by showing that auditory influences are not limited to low-level perceptual events such as flashes and tones (Shams et al., 2000; Koelewijn et al., 2010), but also affect the recognition of symbolic, culturally mediated stimuli such as written words. This aligns with evidence that language perception is intrinsically multimodal, involving close interaction between auditory and visual channels (Sekuler et al., 1997; Spence, 2011; Giraud & Poeppel, 2012).

Multi modal integration has been known from many examples where the perception of a visual stimulus is influenced by varying auditory stimuli (Large & Jones, 1999; Schroeder & Lakatos, 2009). This is the first time, however, that such an integration is relevant for the perception of the written word. This supports the dynamic attending theory (Jones, 1976; Jones and Boltz, 1989), which posits that attention fluctuates rhythmically and can be entrained by external periodic events. The observed persistence of the beat effect after stimulus offset is consistent with the idea of an isochronous oscillatory process that continues to organize attentional resources in line with previously established temporal expectations.

At a mechanistic level, this paradigm highlights how auditory rhythms can serve as temporal scaffolds for visual processing. One possibility is that periodic auditory stimulation sharpens the phase alignment of attentional peaks, thereby increasing the likelihood that visual stimuli presented in phase are encoded. Alternatively, auditory cues may act by reducing uncertainty in the timing of visual input, thereby lowering the cognitive load required for detection.

Distinguishing between these alternatives will require further work, for example by manipulating the predictability or complexity of the rhythmic sequence.

In sum, the present study shows that written word perception, like other domains of sensory processing, is sensitive to rhythmic structure and susceptible to cross-modal entrainment. The results suggest that attentional oscillations constitute a general mechanism by which the brain integrates information across modalities, extending beyond simple sensory events. To conclude, the perception of language is intrinsically multi-modal, even in such culturally created media as the written word.

## Acknowledgements

We thank Nikola Falk for her help testing participants and Jinook Oh for his help in building the program. This research was supported by Austrian Science Fund (FWF) DK Grant “Cognition & Communication” (#W1262-B29).

